# Capacity to erase gene occlusion is a defining feature distinguishing naive from primed pluripotency

**DOI:** 10.1101/2021.05.04.442547

**Authors:** Kara M. Foshay, Jae Hyun Lee, Li Zhang, Croydon J. Fernandes, Bohou Wu, Jedidiah Gaetz, Samuel W. Baker, Timothy J. Looney, Andy Peng Xiang, Guoping Fan, Bruce T. Lahn

## Abstract

Pluripotent stem cells can exist in either the naive state representing a developmental blank slate or the downstream primed state poised for differentiation. Currently, known differences between these two states are mostly phenomenological, and none can adequately explain why the two states should differ in developmental priming. Gene occlusion is a mode of epigenetic inactivation that renders genes unresponsive to their cognate transcriptional activators. It plays a crucial role in lineage restriction. Here, we report that a defining feature distinguishing the two pluripotent states lies in the ability of naive but not primed cells to erase occlusion. This “deocclusion” capacity requires Esrrb, a gene expressed only in the naive but not primed state. Notably, Esrrb silencing in the primed state is itself due to occlusion. Collectively, our data argue that the Esrrb-dependent deocclusion capacity in naive cells is key for sustaining naive pluripotency, and the loss of this capacity in the primed state via the occlusion of Esrrb poises cells for differentiation.

## INTRODUCTION

In mammals, the early embryo harbors uncommitted pluripotent stem cells capable of differentiating into all lineages of the body (De Los Angeles et al. 2015). It is now recognized that pluripotency is not a single state, but a continuum wherein cells become increasingly capacitated to undergo differentiation. This continuum is bracketed on one end by the naive state representing the starting blank slate of development, and on the other end by the primed state that is developmentally further downstream where cells become poised to differentiate (Nichols and Smith 2009; Kinoshita and Smith 2018). Both naive and primed pluripotent cells can contribute to all cell types in the embryo proper and both express canonical pluripotency genes such as Pou5f1 (Oct4), Nanog and Sox2 (Brons et al. 2007; Tesar et al. 2007). These similarities notwithstanding, many differences between them have been reported. Primed cells, in contrast to naive cells, are unresponsive to LIF, cannot contribute to blastocyst stage chimeras, and show the onset of X-inactivation. While transcriptome profiles of the two pluripotent states are highly similar, there are a number of genes displaying strong differential expression, and the two states also show some differences in miRNA profile and enhancer usage (Brons et al. 2007; Tesar et al. 2007; Jouneau et al. 2012; Buecker et al. 2014; Factor et al. 2014). However, these differences are largely phenomenological and none can satisfactorily explain, at a fundamental level, what makes the naive state developmentally blank and the primed state poised for differentiation.

In this study, we examine whether gene occlusion plays a role in the differential developmental priming of these two pluripotent states. Gene occlusion is a mode of epigenetic silencing wherein chromatin-based, cis-acting mechanisms render genes unresponsive to transacting transcriptional activators in the cell (Lee et al. 2009a; Lahn 2011). Opposite the occluded state is the competent state wherein genes are competent to respond to trans-acting factors. Occluded genes are thus stably silenced irrespective of whether their cognate transcriptional activators are present, whereas competent genes can be either active or silent depending on the presence or absence of their activators in the cell.

During lineage differentiation, cells not only gain phenotypic identities of the committed lineage, but they also lose developmental potential for all the other lineages – a process known as lineage restriction. We previously proposed a model of lineage restriction that invokes gene occlusion as a key underlying mechanism (Lahn 2011). The model posits that in pluripotent stem cells, virtually all genes are in the competent state as they are fully responsive to transcriptional activators, but as lineage differentiation proceeds, lineage-inappropriate genes shift irreversibly to the occluded state where they are no longer responsive to activators. This leads to the permanent silencing of these genes, thus restricting the developmental plasticity of cells to only the committed lineage. In support of this model, we uncovered the prevalence of occluded genes in a variety of somatic cell types and demonstrated the importance of occlusion in restricting the identities of these cells (Lee et al. 2009a; Lee et al. 2009b; Gaetz et al. 2012; Looney et al. 2014). We further demonstrated that ESCs, unlike somatic cells, possess a unique ability to erase occlusion across the genome (Foshay et al. 2012). Such “deocclusion” capacity could enable naive pluripotent cells to establish and sustain full competency of the genome before the onset of lineage differentiation.

We hypothesized that primed pluripotency, being a developmental intermediate between the naive state and the differentiated state, might lose the deocclusion capacity as a prerequisite for priming the genome for differentiation and lineage restriction (Lahn 2011). Here, we present evidence in support of this hypothesis. Additionally, we show that Esrrb, a pluripotency gene highly expressed in the naive state but silent in the primed state, is a critical component of the deocclusion machinery in naive cells. Furthermore, we demonstrate that the silencing of Esrrb in primed cells is itself rendered by occlusion, and this leads to the loss of the deocclusion capacity in these cells.

## RESULTS

### Naive and primed states differ in their ability to erase occlusion

Our main assay for identifying occluded genes is cell fusion (Lee et al. 2009a; Looney et al. 2014). If a gene is silent in a given cell type, and this silencing persists even after fusion to another cell type in which the ortholog is active, then this silencing is due to occlusion, for it is unresponsive to transcriptional activators availed by the fusion. But if the silent gene is activated upon fusion, then it is considered “activatable”, for it is responsive to activators. Here, we employed the widely used mouse embryonic stem cell (ESC) line E14 (Doetschman et al. 1987) representing the naive state and the inhouse-derived mouse epiblast stem cell (EpiSC) line G14 representing the primed state in a series of fusion studies to examine gene occlusion in these cells. Transcriptome profiling by RNA-seq confirmed that both E14 and G14 expressed canonical pluripotency genes Pou5f1 and Nanog. Additionally, strong differential expression was confirmed for known naive-specific genes such as Esrrb and Klf4 as well as known primed-specific genes such as Fgf5 (Table S1). Following similar approaches as our previous studies (Lee et al. 2009a; Foshay et al. 2012; Looney et al. 2014), we set out to fuse G14 with somatic cells of rat origin in order to identify genes active in EpiSCs but occluded in the somatic cell type. After attempting to fuse G14 with a number of rat somatic cell lines, we found that the rat neuroblastoma cell line B35 could yield a sufficient number of viable hybrid cells for analysis. We therefore primarily used B35 for subsequent fusion studies.

After fusing G14 and B35, hybrid cells were maintained under EpiSC culture condition where they displayed similar morphology as unfused EpiSCs. Our previous fusion studies showed that by day 8 post fusion, transcription factors would have had ample time to act on their target genes (Looney et al. 2014). At this point, for genes showing strong differential expression between the two genomes of the hybrid cell, the silent copies are considered to be occluded. We therefore collected populations of hybrid cells at day 8, referred to as G14-B35(d8). To examine the effect of fusion at the clonal level, we also picked and expanded two hybrid clones, named G14-B35(clone9) and G14-B35(clone11). Transcriptome profiling of the hybrids as well as the unfused parental cells uncovered a list of robustly occluded genes in B35 (Table S2). These genes showed greater than ten-fold expression in unfused G14 relative to their orthologs in unfused B35, with G14 expression reaching a minimum of 10 transcripts per genome. In the G14-B35(d8) hybrid, the G14 and B35 copies of these genes continued to show greater than ten-fold expression difference (Fig 1A, top row). In the two hybrid clones, strong differential expression persisted despite the fact these cells that carry both genomes have been actively proliferating for several months prior to transcriptome analysis.

**Figure 1.**
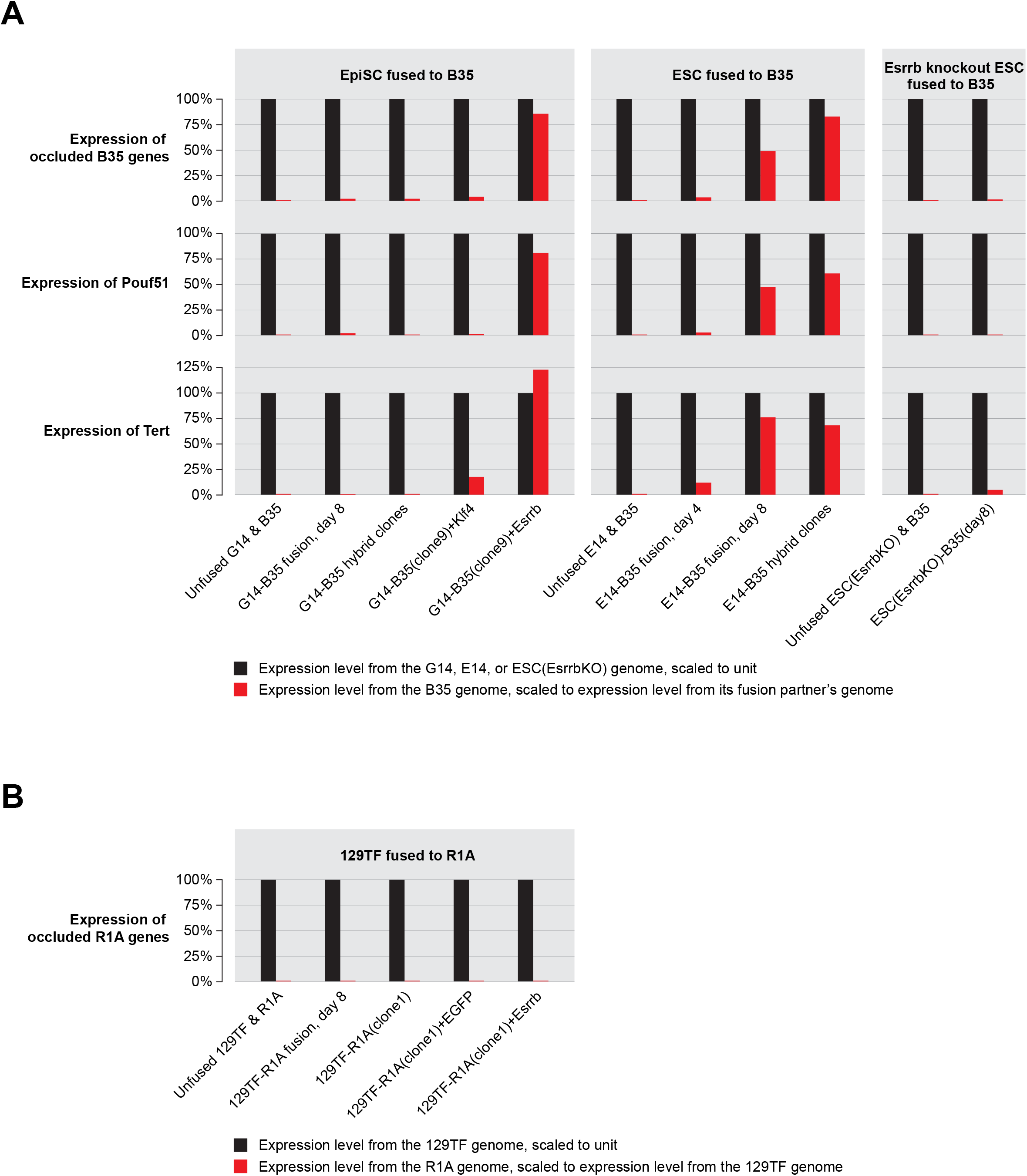
Expression of occluded genes in various cell fusion contexts. **(A)** Expression of occluded B35 genes in fusions between B35 and various pluripotent cell types. The left column shows fusions between the EpiSC cell line G14 and B35, including (from left to right): unfused G14 and B35, G14-B35 fusion population sample at day 8, average of two G14-B35 hybrid clones (#9 and 11), G14-B35(clone9) hybrid with ectopic Klf4 expression, and G14-B35(clone9) hybrid with ectopic Esrrb expression. The middle column shows fusions between the ESC cell line E14 and B35, including: unfused E14 and B35, E14-B35 fusion population sample at day 4, E14-B35 fusion population at day 8, and average of three E14-B35 hybrid clones. The right column shows fusions between the Esrrb knockout ESC line ESC(EsrrbKO) and B35, including: unfused samples and the fusion population sample at day 8. The top row shows the average expression of the occluded B35 genes from the B35 genome and that of their orthologs in the fusion partner’s genome. The middle and bottom rows show expression of Pou5f1 and Tert, respectively. **(B)** Expression of occluded R1A genes in 129TF-R1A fusions. Samples include (from left to right): unfused 129TF and R1A, 129TF-R1A fusion population at day 8, 129TF-R1A(clone1) hybrid, 129TF-R1A(clone1) hybrid with ectopic EGFP expression, 129TF-R1A(clone1) hybrid with ectopic Esrrb expression.

We next fused the ESC cell line E14 to B35 to examine how the occluded B35 genes identified in the G14-B35 fusion would behave in this fusion. Fused cells were maintained under ESC culture conditions where they assumed ESC morphology. Populations of fused cells were collected at day 4 and 8 post fusion, referred to as E14-B35(d4) and E14-B35(d8), respectively. Additionally, three hybrid clones were isolated and expanded, named E14-B35(clone1-3). RNA-seq on these samples showed that the occluded B35 genes identified in the G14-B35 fusion were activated in the E14-B35 fusion, indicating that their occluded status was erased (Fig 1A, top row). This is consistent with ESCs possessing a deocclusion machinery as reported previously (Foshay et al. 2012). Informatively, there is barely any activation of these genes in E14-B35(d4) but robust activation in E14-B35(d8), with even greater activation in the E14-B35 hybrid clones. This slow kinetics in the activation of occluded B35 genes is in line with our previous study showing that the deocclusion machinery in ESCs requires cell division to exert its effect (Foshay et al. 2012).

It is important to note that one of the genes on the list of occluded B35 genes is the core pluripotency factor Pou5f1 (Fig 1A, middle row). It is abundantly expressed in both ESCs and EpiSCs but silent in all somatic cell types examined including B35. In the G14-B35 fusion, both the d8 population sample and the hybrid clones showed high levels of Pou5f1 expression from the G14 genome but minimal expression from the B35 genome. In contrast, in the E14-B35 fusion, the d8 population sample and the hybrid clones showed high levels of Pou5f1 expression from both E14 and B35 genomes.

Another noteworthy gene on the list is the telomerase gene Tert (Fig 1A, bottom row). Tert is expressed in embryonic stem cells, germ cells, and many types of cancer cells, and is silent in most differentiated somatic cell types (Flores et al. 2006). We have previously found that the silencing of Tert in somatic cells is mediated by occlusion (unpublished data). In the cell lines employed in this study, Tert is expressed at moderate levels in G14 and E14, but silent in B35. In the G14-B35 fusion, both d8 population sample and the hybrid clones showed moderate Tert expression from the G14 genome but little expression from the B35 genome, indicating that Tert is occluded in B35. In the E14-B35 fusion, by contrast, both the d8 population sample as well as the hybrid clones showed comparable levels of Tert expression from both E14 and B35 genomes, indicating that Tert in the B35 genome is deoccluded upon fusion to E14.

Together, these data reveal a striking distinction between ESCs and EpiSCs, namely, the presence of a deocclusion machinery in ESCs that is absent in EpiSCs. In the context of fusion to somatic cells, this deocclusion machinery enables ESCs but not EpiSCs to reprogram and activate occluded genes in the somatic genome, with Pou5f1 and Tert being notable examples.

### Naive and primed states differ in their ability to remodel chromatin of occluded genes

Next, we examined whether the differential ability of EpiSCs and ESCs to erase occlusion extended to the remodeling of chromatin. We mapped chromatin openness of the occluded genes describe in the preceding section around their transcription start sites (TSSs) by DNase-seq in five samples: G14, E14, B35, G14-B35(clone9), and E14-B35(clone2). In unfused G14, E14 and B35, these genes displayed DNase hypersensitivity profiles consistent with their expression patterns, namely, there is robust DNase signal in both G14 and E14 but little signal in B35 (Fig 2A&B, left panels). In the G14-B35(clone9) hybrid, the TSSs of the B35 copies of the genes showed only slight increase in DNase hypersensitivity signal compared to unfused B35, whereas the G14 copies of the genes showed similarly strong signal as in unfused G14 (Fig 2A, right panel), indicating deficient chromatin remodeling. In contrast, in the E14-B35(clone2) hybrid, strong DNase signal is seen for the TSSs of both E14 and B35 copies of these genes (Fig 2B, right panel), indicating that the B35 copies of the genes have been effectively remodeled from closed to open chromatin in the hybrid. When examined on a gene by gene basis, similar patterns were observed (Fig 2C-F).

**Figure 2.**
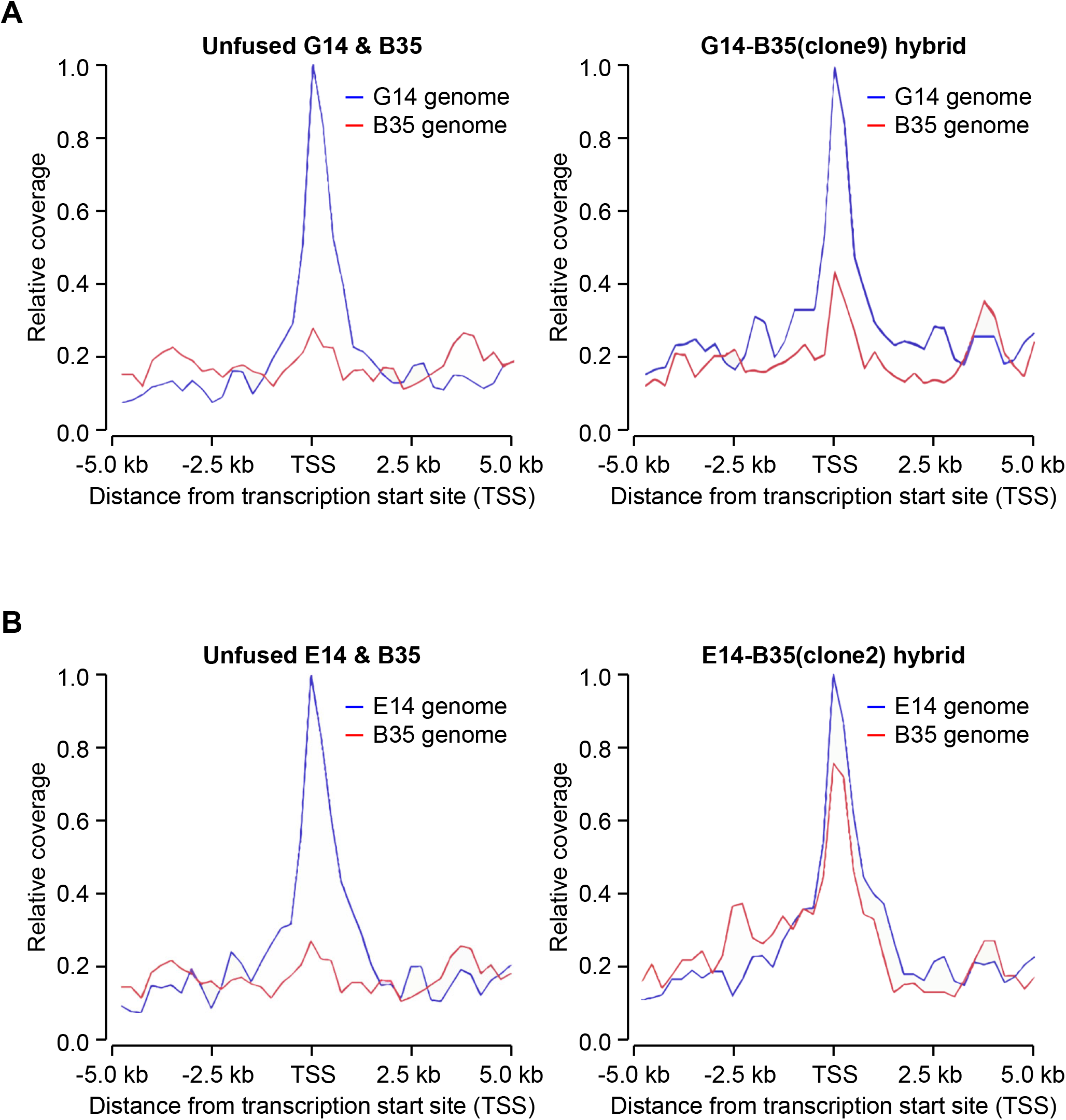
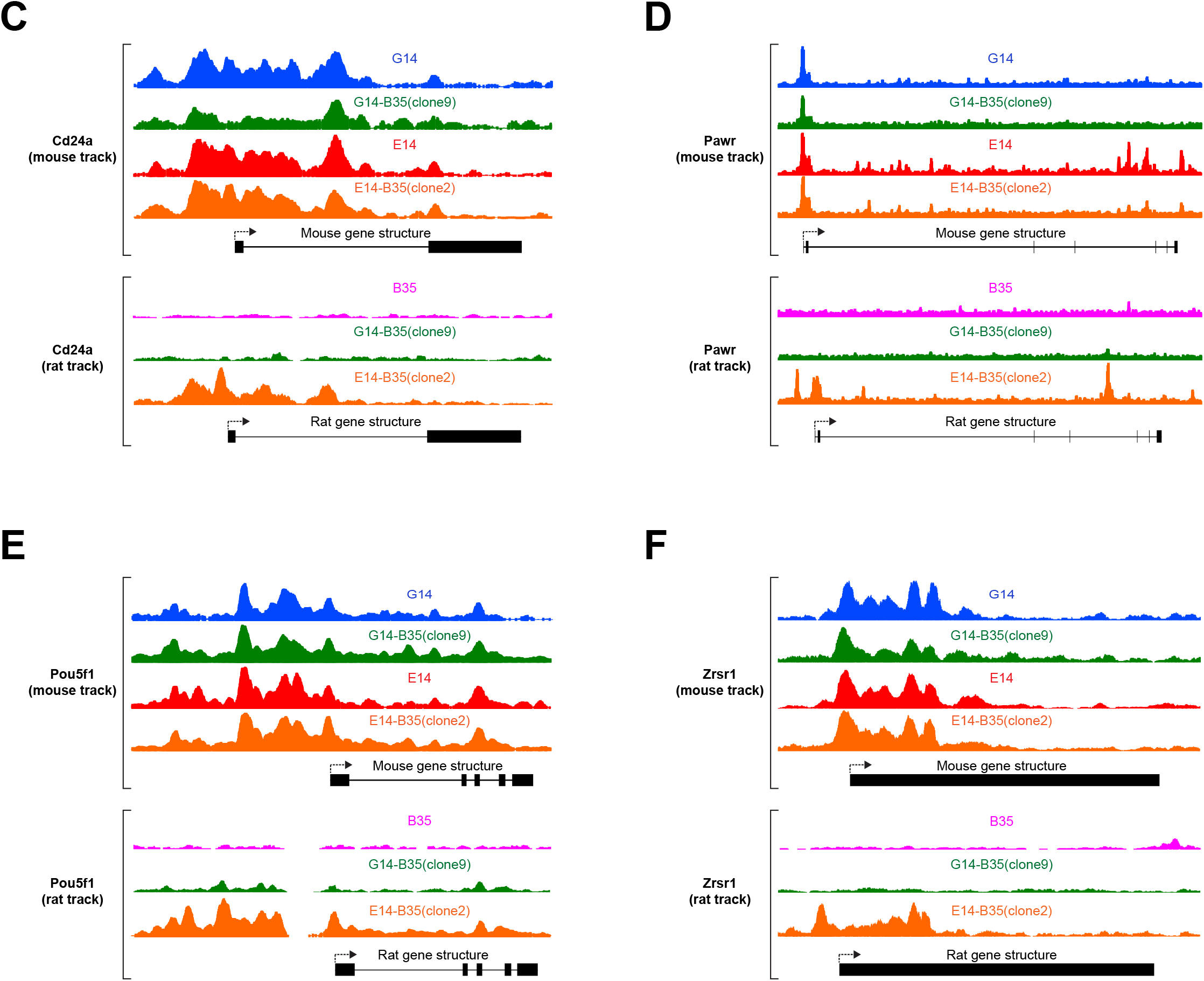
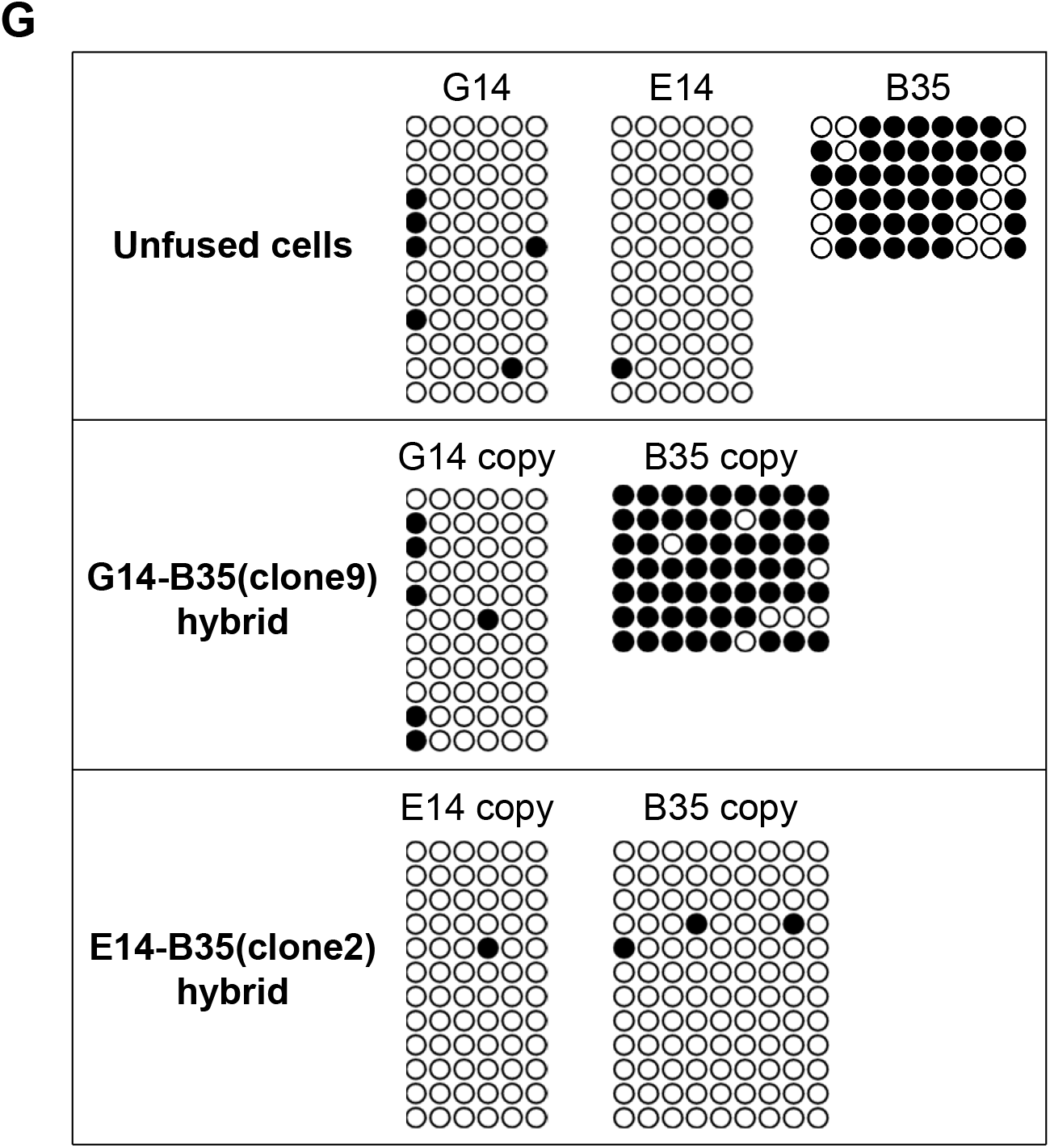
Differential ability of ESCs and EpiSCs to erase occlusion extends to the remodeling of chromatin. **(A-C)** DNase-seq was used to map chromatin openness in the EpiSC cell line G14, the ESC cell line E14, the somatic cell line B35, G14-B35(clone9) hybrid, and E14-B35(clone2) hybrid. **(A)** Average DNase hypersensitivity profiles of the occluded B35 genes (red line) and their orthologs in the G14 genome (blue line) across 10 kb centered around transcription start site (TSS). The left panel shows data from unfused G14 and B35; the right panel shows data from G14-B35(clone9) hybrid. **(B)** Similar analysis as in (A) except E14 is used in the experiments instead of G14. The left panel shows data from unfused E14 and B35, while the right panel shows data from E14-B35(clone2) hybrid. **(C-F)** Representative DNase hypersensitivity profiles of individual genes. **(G)** Analysis of Pou5f1 distal enhancer CpG methylation in G14, E14, B35, G14-B35(clone9) hybrid, and E14-B35(clone2) hybrid. Each column of circles represents a CpG site while each row represents a sequenced clone covering all the sites (see Methods). Black and white circles represent, respectively, methylated and unmethylated CpGs.

The CpG methylation status of Pou5f1 cis-regulatory region has been shown to be a marker of pluripotency, with pluripotent cells showing low methylation while somatic cells showing high methylation (Han et al. 2011; Choi et al. 2016). We examined Pou5f1 distal enhancer methylation in the above samples. Unfused G14 and E14 both showed little methylation whereas B35 showed strong methylation (Fig 2G, top). In G14-B35(clone9) hybrid, little methylation was seen in the G14 copy of Pou5f1 just like in unfused G14 whereas heavy methylation was present in the B35 copy, indicating a failure of the hybrid to reprogram the methylation status of the B35 copy (Fig 2G, middle). In contrast, in E14-B35(clone2) hybrid, both E14 and B35 copies are devoid of methylation, indicating full reprograming of the methylation status of the B35 copy (Fig 2G, bottom).

Collectively, the above data show that the differential ability of naive and primed states in erasing occlusion extends to the chromatin level, with the naive but not the primed state capable of remodeling the chromatin of occluded genes to an open configuration.

### Ectopic Esrrb expression promotes deocclusion in primed pluripotent cells

While naive and primed pluripotent states share similar gene expression patterns, there exist a number of genes that are expressed in the naive state but silent in the primed state. We speculated that some of these differentially expressed genes may encode critical components of the deocclusion machinery present in the naive state, and that its silencing in the primed state is responsible for the loss of the deocclusion capability in the primed state.

We focused on two candidate genes, Klf4 and Esrrb, both are expressed in the naive but silent in the primed state (Brons et al. 2007; Tesar et al. 2007), and are implicated in promoting naive pluripotency (Feng et al. 2009; Guo et al. 2009; Buganim et al. 2014). We transduced the G14-B35(clone9) hybrid with lentivirus expressing either Klf4 or Esrrb. After extended culture, cells were harvested and subjected to RNA-seq. In the Klf4-transduced sample, the occluded B35 genes remained silent in the B35 genome whereas their G14 orthologs in the same fused cells stayed active, indicating that the occluded status of these genes was not significantly reprogramed (Fig 1A). By contrast, in the Esrrb-transduced sample, the occluded B35 genes were robustly activated, reaching expression levels comparable to their G14 orthologs in fused cells (Fig 1A). Thus, expression of ectopic Esrrb but not Klf4 is sufficient to confer deocclusion capability to primed pluripotent cells that otherwise lack this capability.

### Esrrb is necessary for the deocclusion capacity in naive pluripotent cells

We next examined whether Esrrb is necessary for the deocclusion capability present in the naive pluripotent state. We utilized an Esrrb knockout ESC line in which exon 2 is deleted on both copies of the gene, referred to hereon as ESC(EsrrbKO) (Martello et al. 2012). RNA-seq analysis confirmed that this cell line has a characteristic ESC transcriptome profile (Table S1). ESC(EsrrbKO) was fused with B35, cultured for 8 days and harvested for transcriptome profiling. For the occluded B35 genes, their behavior in this fusion is very similar to that of the G14-B35 fusion, namely the ESC(EsrrbKO) copies of these genes and the B35 copies maintained strong differential expression in hybrid cells just as in unfused parental cells (Fig 1A). This stands in sharp contrast to the fusion between B35 and the wildtype ESC line E14, where the occluded B35 genes became strongly activated in the hybrid (Fig 1A). This result shows that Esrrb is necessary for the deocclusion capability present in naive pluripotent cells. Combining this finding with that of previous sections, we conclude that the deocclusion capacity in the naive pluripotent state is lost when cells transition into the primed state, and that this loss is rendered through the silencing of Esrrb in the primed state.

### Esrrb is occluded in primed pluripotent cells

We hypothesized that the silencing of Esrrb in primed pluripotent cells is itself the result of occlusion. One apparent strategy to test this is to fuse ESCs with EpiSCs, and examine whether the EpiSC copies of Esrrb in hybrid cells would stay silent whereas the ESC copies would stay active. However, there are two problems with this approach. First, because ESCs possess the deocclusion capacity, any occluded genes in EpiSC would have their occlusion erased upon fusion to ESCs. Second, if we were to fuse wildtype ESCs and EpiSCs, both of mouse origin, there would be no easy way to tell if Esrrb transcripts in the hybrid are produced by ESC or EpiSC copies of the gene. Conveniently, both difficulties can be overcome by using Esrrb knockout ESCs in the fusion. As shown in the preceding section, Esrrb knockout abolishes the deocclusion capacity in ESCs. Furthermore, the Esrrb knockout allele lacks exon 2 but retains other exons. It can therefore still undergo transcription except its mRNA lacks exon 2. This would allow the mapping of Esrrb transcript in hybrid cells specifically to either the Esrrb knockout ESC genome or the EpiSC genome based on the presence or absence of exon 2. We fused ESC(EsrrbKO) to G14, and performed RT-PCR on fused cells harvested 16 days after fusion. Two amplicons spanning exons 2-4 of Esrrb, which are designed to only amplify the wildtype allele but not the knockout allele, did not produce any RT-PCR signal from hybrid cells (Fig 3A). This indicates that the wildtype Esrrb in the G14 genome is silent in the hybrid. By contrast, two other amplicons, one spanning exons 3-5 of Esrrb and the other exons 4-6, which can amplify both the wildtype allele and the knockout allele, produced RT-PCR signal from fused cells (Fig 3A). In light of the fact that wildtype Esrrb in the G14 genome of hybrid cells is silent, this signal must come from the ESC(EsrrbKO) genome. Thus, Esrrb is differentially expressed in the hybrid, being active from the ESC(EsrrbKO) genome but silent from the G14 genome, indicating the occluded status of the gene in G14.

**Figure 3.**
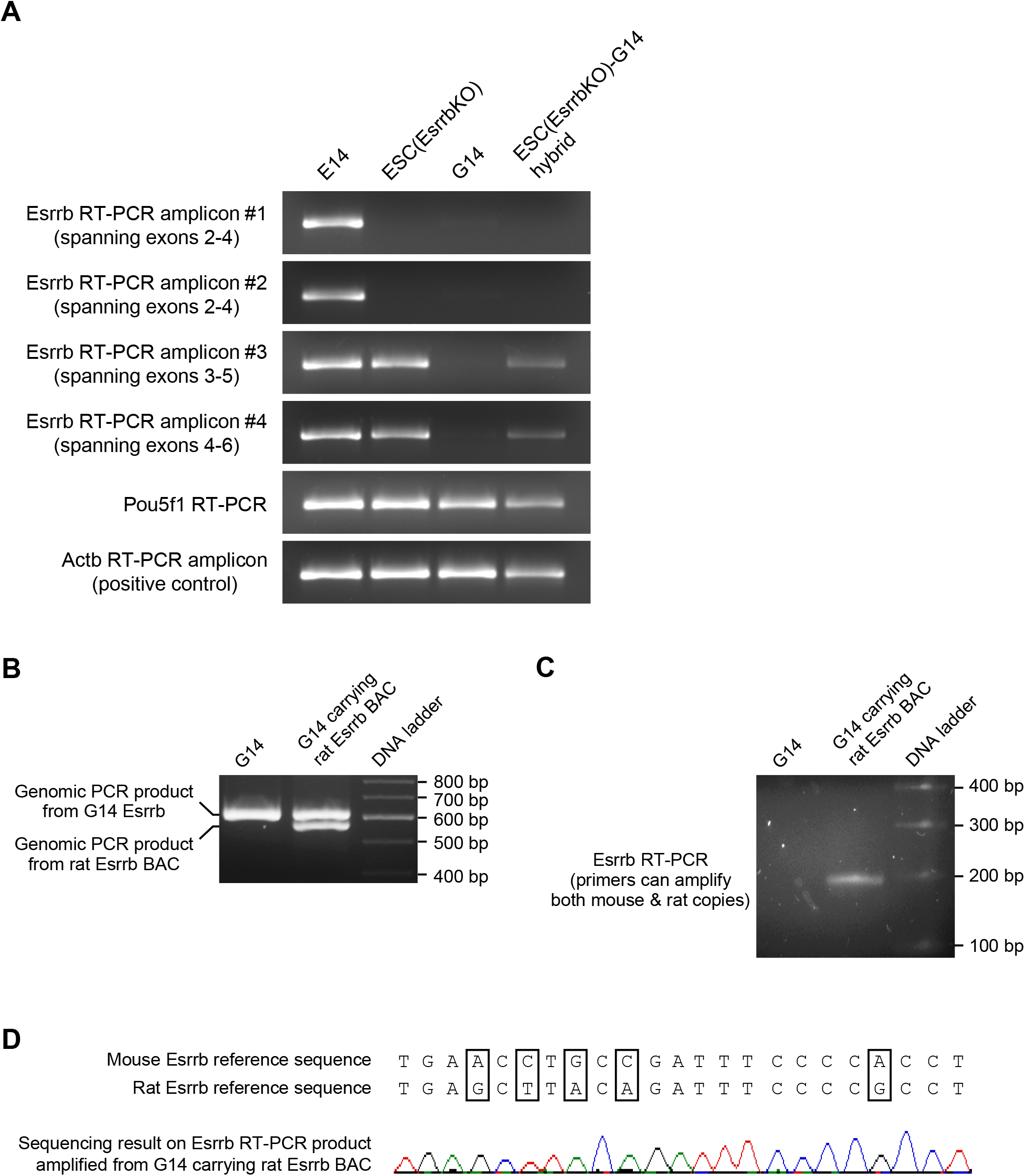
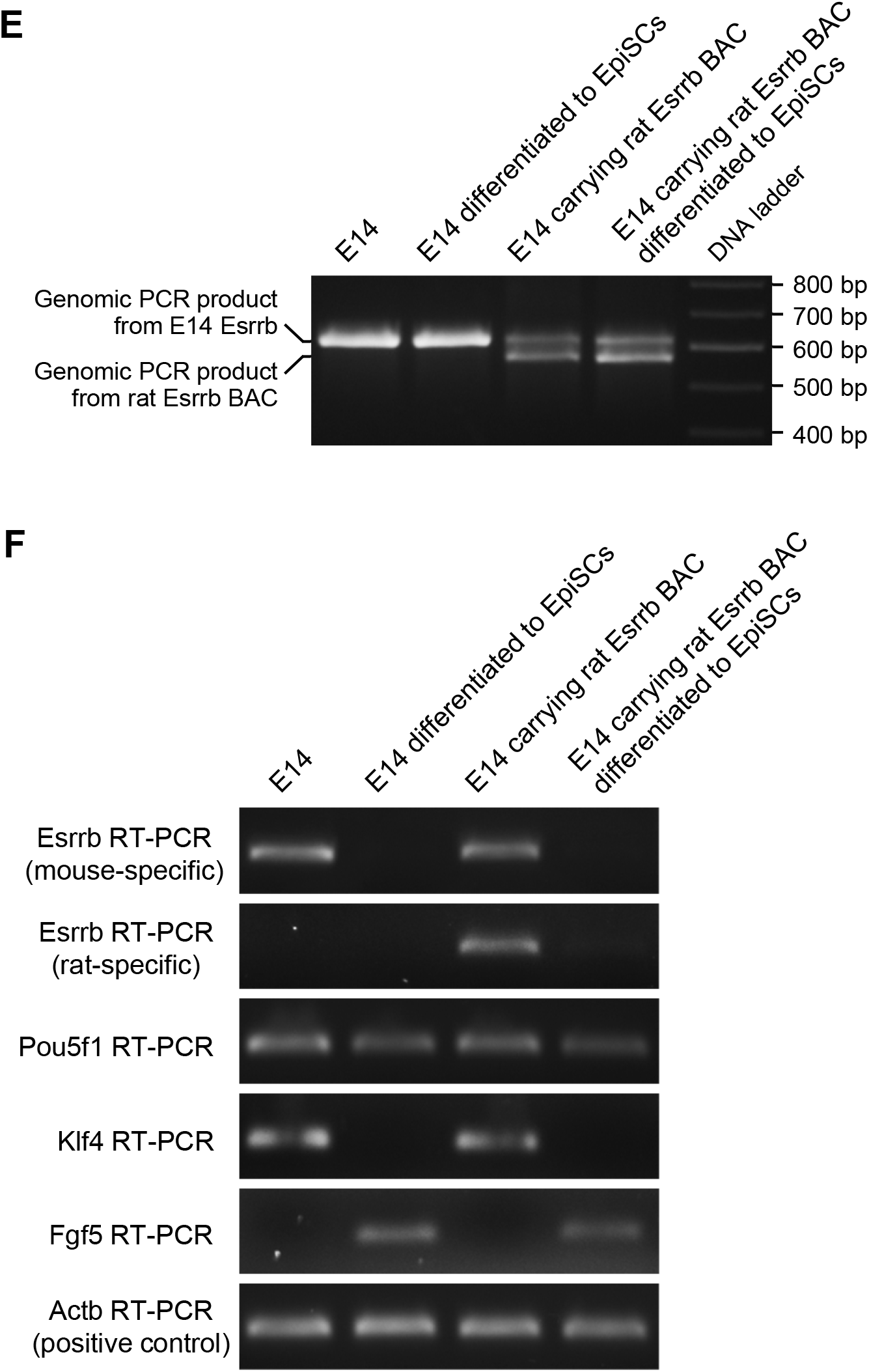
Occlusion of Esrrb in EpiSCs. **(A)** Cell fusion assay showing that Esrrb is occluded in EpiSCs. Esrrb Expression was interrogated by four RT-PCR amplicons, each in four samples including (from left to right): the ESC cell line E14, the Esrrb knockout ESC cell line ESC(EsrrbKO), the EpiSC cell line G14, and ESC(EsrrbKO)-G14 hybrid. The top two Esrrb amplicons were designed to only amplify the wild type Esrrb transcript, whereas the bottom two Esrrb amplicons were designed to amplify transcript from both the wild type and the knockout Esrrb alleles. Pou5f1 and Actb amplicons were included as controls. The results showed that in ESC(EsrrbKO)-G14 hybrid cells, Esrrb is expressed from the ESC(EsrrbKO) genome but not the G14 genome, indicating that it is occluded in G14. **(B, C & D)** BAC transgene assay showing that Esrrb is occluded in the EpiSC cell line G14. A BAC carrying the entire rat Esrrb gene was integrated into the G14 genome by piggyBac-mediated transposition, which was confirmed by genomic PCR using primers that can amplify both the mouse endogenous Esrrb and the rat Esrrb BAC but producing different-size products (B). Using RT-PCR primers designed to amplify both G14 (mouse) and BAC (rat) Esrrb transcript, Esrrb expression was detected in the G14 cells carrying the rat Esrrb BAC but not the wildtype G14 (C). Sequencing of the RT-PCR product spanning polymorphic bases between mouse and rat versions of Esrrb (boxed) confirmed that expression originated from the BAC copy of the Esrrb (D). **(E & F)** Esrrb occlusion is established by a machinery that functions transiently during the naïve-to-primed transition. The same rat Esrrb BAC as used in panel B was integrated into the E14 genome by piggyBac-mediated transposition, which was confirmed by the same genomic PCR as used in panel B (E). Transgene-positive cells were then differentiated into EpiSCs. RT-PCR using primers specific to the mouse endogenous Esrrb or the rat Esrrb transgene showed that they both became silenced in the EpiSCs differentiated from E14 (F). Mouse Pou5f1, Klf4, Fgf5 and Actb were also analyzed to confirm the naïve and primed states of the cells before and after differentiation, respectively (F).

A second assay we developed to identify occluded genes involves the introduction of BAC transgenes into cells (Gaetz et al. 2012). BACs are typically large enough to accommodate all cis-regulatory regions of a gene, and the lack of any initial chromatin modifications on BAC DNA ensures that BAC transgenes are not occluded when first entering the cell (Gaetz et al. 2012). As such, the expression status of the BAC transgene provides a readout of the transacting milieu of the cell. Active expression would indicate that the cell contains transcriptional activators needed to drive expression of the gene. If the BAC transgene is expressed while the endogenous ortholog remains silent, then the silencing of the endogenous gene is necessarily due to occlusion (Gaetz et al. 2012). Here, we introduced a BAC containing the entire rat Esrrb gene into G14 cells via piggyBac transposition. Transgene-positive cells were selected and cultured for an extended period. Genomic PCR confirmed the presence of both rat and mouse copies of Esrrb in the cells (Fig 3B). In contrast, RT-PCR showed that the rat Esrrb transgene but not the endogenous Esrrb was expressed in these cells (Fig 3C, D). This result again indicates the occluded status of endogenous Esrrb in G14. More importantly, it shows that the cellular milieu of G14 contains all the trans-acting factors necessary to drive Esrrb transcription, as evidenced by the expression of the Esrrb BAC transgene. Thus, it can be concluded that the endogenous Esrrb in G14 is rendered silent by occlusion in a cellular milieu that is otherwise supportive of its expression.

### Esrrb occlusion is established by a machinery that functions transiently during the naive-to-primed transition

The fact that the rat Esrrb BAC transgene introduced into EpiSCs does not itself undergo occlusion suggests that the occlusion of the endogenous Esrrb is established by a machinery that functions transiently during the naive-to-primed state but not after the transition. To further test this idea, we introduced the same rat Esrrb BAC into E14 by piggyBac transposition, and differentiated the cells into EpiSCs. Genomic PCR confirmed the successful integration of the BAC into the genome (Fig 3E). RT-PCR expression analysis of Pou5f1, Klf4 (a naive-specific marker), and Fgf5 (a primed-specific marker) confirmed the naive and primed states of the cells before and after differentiation, respectively (Fig 3F). Importantly, both the endogenous Esrrb and rat Esrrb transgene became silenced in the EpiSCs differentiated from E14 (Fig 3F), which stands in contrast to the active expression of the rat Esrrb transgene introduced directly into the wildtype EpiSC line G14. This result supports the notion that the machinery responsible for establishing the occluded status of Esrrb functions transiently during the naive-to-primed transition but not after the cells have fully settled in the primed state.

### Esrrb is occluded in somatic cells

Esrrb is silent in a variety of somatic cell lines that we studied. We wished to examine whether this is also due to occlusion as is the case in EpiSCs. Our previous study found that in ESC-somatic fusions, occluded genes in the somatic genome showed a characteristically slow activation kinetics whereas activatable genes in the somatic genome displayed rapid activation in the same fusions (Foshay et al. 2012). We therefore examined the activation kinetics of somatic copies of Esrrb in the E14-B35 hybrids as well as in the previously reported fusion between E14 and the rat fibroblast line R1A (Foshay et al. 2012). In both cases, somatic copies of Esrrb showed slow activation characteristic of occluded genes in the somatic genome, consistent with Esrrb being occluded in B35 and R1A (Fig 4A).

**Figure 4.**
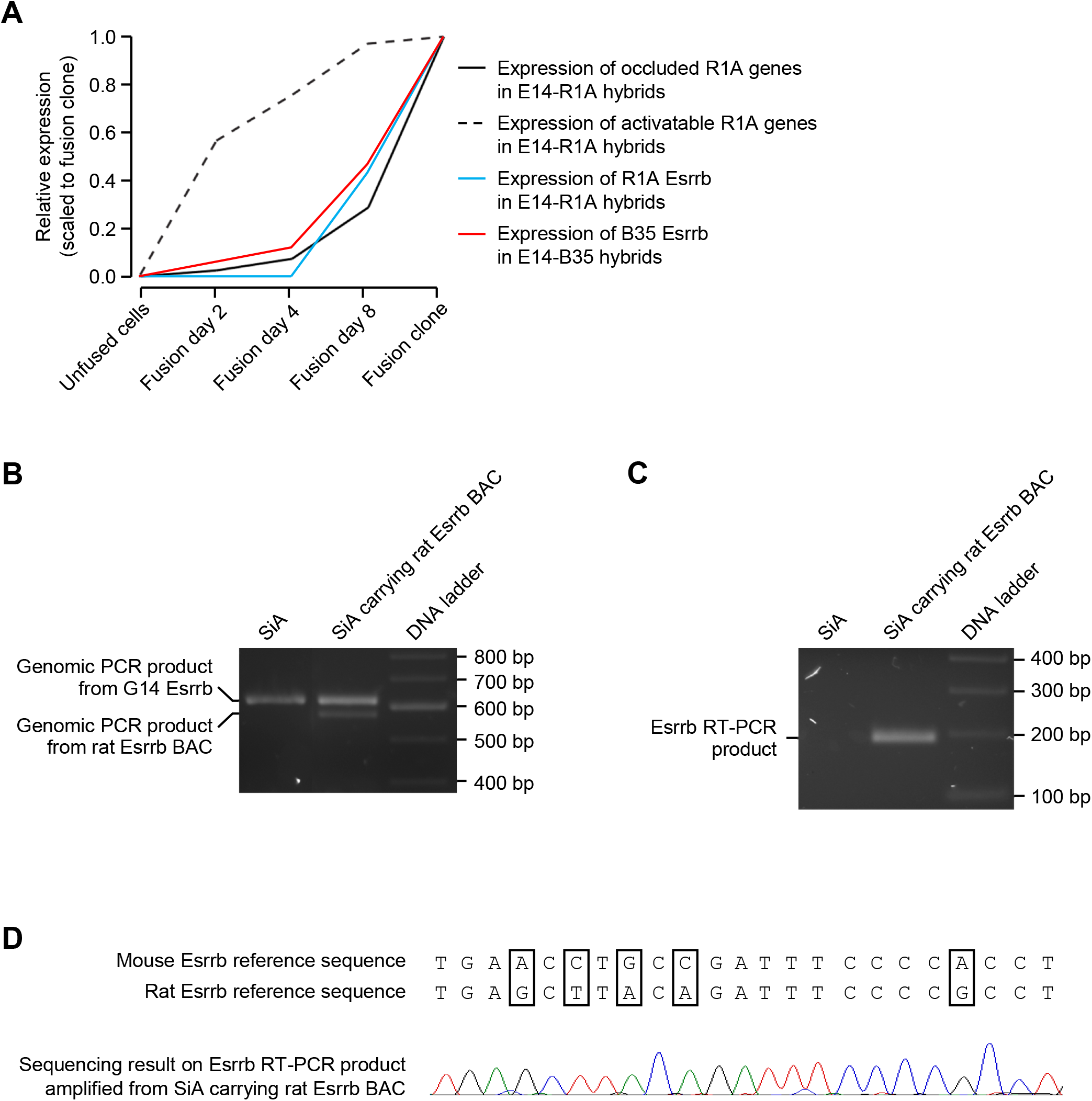
Evidence that Esrrb is occluded in somatic cells. **(A)** Occluded status of Esrrb in somatic cells revealed by its slow activation kinetics in ESC-somatic fusions. In fusions between the mouse ESC line E14 and the rat cell line R1A, activatable R1A genes showed rapid activation whereas occluded R1A genes showed slow activation (Foshay et al., 2012). Esrrb in rat cell lines R1A or B35 showed slow activation when these cells were fused to E14, consistent with its occluded status. **(B, C & D)** BAC transgene assay showing that Esrrb is occluded in the somatic cell line SiA. A BAC carrying the entire rat Esrrb gene was integrated into the SiA genome by piggyBac-mediated transposition, which is confirmed by genome PCR (B). Using RT-PCR primers designed to amplify both SiA (mouse) and BAC (rat) Esrrb transcript, Esrrb expression was detected in the SiA cells carrying the rat Esrrb BAC but not the wildtype SiA (C). Sequencing of the RT-PCR product spanning polymorphic bases between mouse and rat versions of Esrrb (boxed) confirmed that expression originated from the BAC copy of the Esrrb (D).

We next used the BAC transgene assay to further confirm the occlusion status of Esrrb in the mouse tail fibroblast line SiA. The same rat Esrrb BAC as used in Figure 3 was introduced into SiA cells by piggyBac transposition followed by extended culture. Genomic PCR confirmed that both rat and mouse copies of Esrrb are present in the cells (Fig 4B). In contrast, RT-PCR showed that only the rat BAC copy of Esrrb was expressed in these cells, demonstrating that endogenous Esrrb is occluded in SiA (Fig 4C, D). This result also indicates that, as in EpiSCs, the cellular milieu of SiA contains all the trans-acting factors necessary to drive Esrrb expression such that the silencing of Esrrb in these cells is solely the result of occlusion and not due to a lack of transcriptional activators.

### Deocclusion requires additional factors besides Esrrb

That ectopic Esrrb expression in EpiSCs is sufficient to impart deocclusion capability to these cells prompted us to examine whether this is also true for somatic cells. We used lentivirus to express ectopic Esrrb in 129TF-R1A(clone1), which is a clonal hybrid line derived from fusion between the mouse cell line 129TF and the rat cell line R1A (Looney et al. 2014). Cells positively transduced with Esrrb lentivirus were selected and cultured for an extended period. In 129TF-R1A(clone1) hybrid transduced with a control lentivirus expressing mCherry, a list of occluded genes in 129TF were identified based on their being robustly expressed in the R1A genome but silent in the 129TF genome (Table S3). In 129TF-R1A(clone1) hybrid cells transduced with the Esrrb lentivirus, these genes remained highly differentially expressed just as in control cells (Fig 1B, Table S3). This result indicates that unlike in EpiSCs, ectopic Esrrb expression in somatic cells is not sufficient to confer deocclusion capability to these cells. Thus, the deocclusion machinery likely requires additional factors present in EpiSCs but not in somatic cells to be fully functional.

### Esrrb binding in ESCs is enriched in genes capable of becoming occluded in later lineage differentiation

We hypothesized that in naive pluripotent cells, the Esrrb protein may preferentially bind to lineage-specific genes capable of becoming occluded in later lineage differentiation to prevent them from becoming prematurely occluded. To test this, we examined published Esrrb ChIP-seq data from mouse ESCs (Chen et al. 2008; Marson et al. 2008). We focused on genes found in our previous study to be either occluded or activatable in somatic cells (Looney et al. 2014), excluding genes actively expressed in ESCs to remove the confounding influence that Esrrb binding is enriched in active genes in ESCs (Table S4 & S5). These genes can be referred to as “prospective” occluded or activatable genes because their occluded or activatable status is manifested in later somatic lineages. For each gene, we examined Esrrb binding from 10 kb upstream of TSS to the end of the gene body. We found that in ESCs, Esrrb binding is significantly enriched in prospective occluded genes relative to prospective activatable genes, with ChIP-seq peaks found in 57% of the former and 32% of the latter (p < 0.00066) (Table S4 & S5). We also examined the binding of the Pou5f1 protein on the same set of genes in ESCs. Overall, Pou5f1 is bound to a smaller fraction of genes as compared to Esrrb. It also shows an enrichment in occluded genes over activatable genes, but to a much lesser extent (29% versus 23% respectively) that is not statistically significant (p < 0.24) (Table S4 & S5). Interestingly, among the Pou5f1-bound prospective occluded genes, the great majority (81%) are also cobound by Esrrb, which is a statistically significant coincidence (p < 0.02) (Fig S1A). Of the cobound genes, most have both Esrrb and Pou5f1 peaks within a 1 kb window, suggesting functional interactions between the two factors on these genes (Table S4 & S5). This pattern is not observed for activatable genes (Fig S1B). The above observation is consistent with the notion that Esrrb, perhaps in conjunction with other factors, act in naive pluripotent cells as defenders against premature occlusion of those genes that are capable of becoming occluded in later somatic differentiation.

## DISCUSSION

In this study, we demonstrated that the primed pluripotent state is more poised for differentiation relative to the naive state because it has lost the capacity to reprogram occluded genes as compared to the naive state. Moreover, our study revealed a series of critical early developmental events leading to lineage restriction (Fig 5). At the start of development, naive pluripotent cells possess a deocclusion machinery that enforces the global competency of the genome. When the naive state transitions into the primed state, cells inherit a competent genome but lose the capacity for deocclusion due to the silencing of Esrrb, a key component of the deocclusion machinery. Notably, Esrrb silencing is itself rendered by occlusion, and furthermore, the occlusion of Esrrb is established by a mechanism that functions only transiently during the naive-to-primed transition. Crucially from this point onward, cells are poised for differentiation, as different sets of gene can now become occluded in different lineages in the absence of the deocclusion machinery. As a result of the occlusion of lineage-inappropriate genes over the course of differentiation, the genomes of differentiated cells lose competency for alternative cell fates and become restricted to their committed lineages.

**Figure 5.**
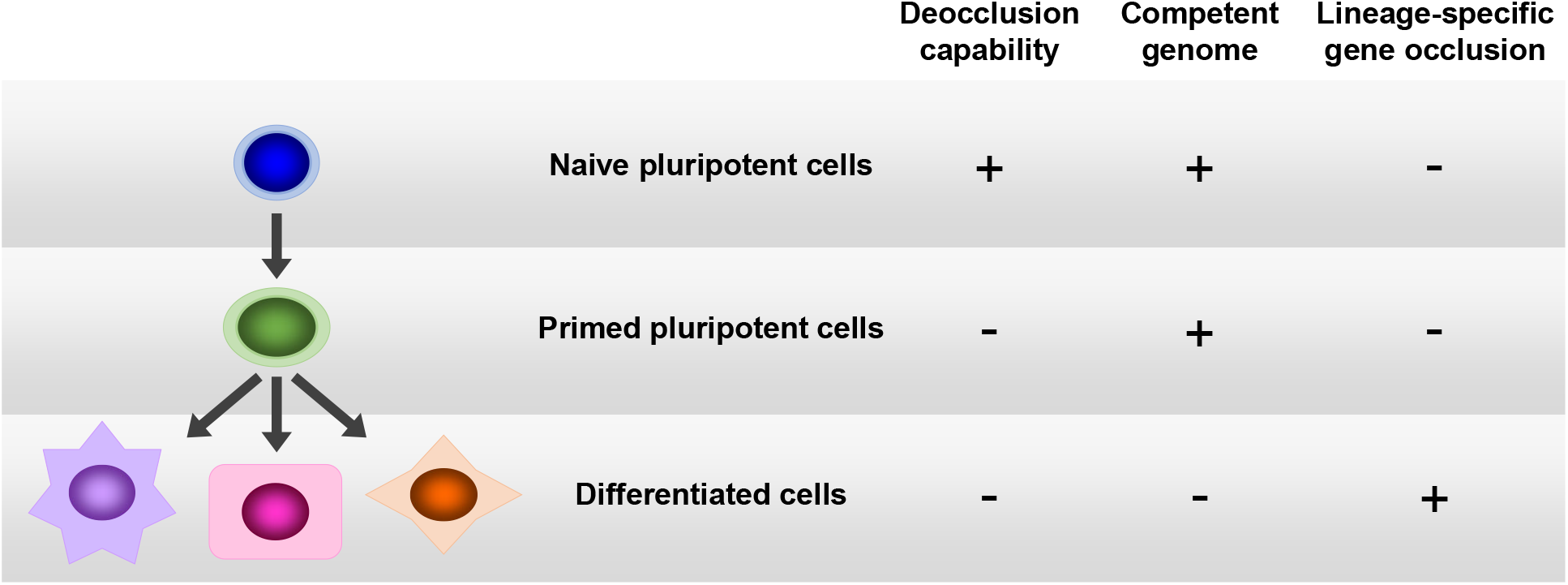
Critical early events leading to lineage restriction. Naïve pluripotent cells are characterized by the presence of the deocclusion capability and a competent genome, and the absence of lineage-specific gene occlusion. When naïve cells transition into primed cells, the deocclusion capability is lost. When primed cells further differentiate into specific lineages, occlusion would permanently shut down the transcriptional potential of specific sets of genes as a means of lineage restriction.

The finding that Esrrb is a critical component of the deocclusion machinery is consistent with previous studies demonstrating a role of Esrrb in promoting pluripotency (Ivanova et al. 2006; van den Berg et al. 2008; Feng et al. 2009; Festuccia et al. 2012; Martello et al. 2012; Papp and Plath 2012; Percharde et al. 2012; Uranishi et al. 2013; Buganim et al. 2014;

Festuccia et al. 2016; Chronis et al. 2017; Adachi et al. 2018; Festuccia et al. 2018a; Festuccia et al. 2018b; Huang et al. 2018; Festuccia et al. 2019). However, Esrrb is not the only component of the deocclusion machinery, as it alone cannot erase occlusion in somatic cells.

Our previous studies have revealed a crucial role of gene occlusion in establishing and restricting the transcriptional identities of differentiated cells. The current study further highlights a key function of occlusion in the regulation of pluripotency even prior to lineage differentiation. The stage is set for future studies to elucidate the molecular details of the machineries that establish, maintain, and erase occlusion in various pluripotent and differentiated cell contexts.

## MATERIALS AND METHODS

### Cell culture and fusion

The mouse cell lines E14 and 129TF, rat cell lines B35 and R1A, and hybrid cell line 129TF-R1A(clone1) were handled as previously described (Foshay et al. 2012; Looney et al. 2014). SiA is the clonal parental line of 129TF. The mouse EpiSC cell line G14 (aka 1117E3) was derived from an E5.5 F1 embryo of a cross between C57BL/6 male and 129Jae female using published method and culture conditions (Brons et al. 2007; Tesar et al. 2007). The Esrrb knockout ES cell line ESC(EsrrbKO) was published previously (Martello et al. 2012).

For E14-B35 and G14-B35 fusions, E14 and G14 cells were first transduced with a lentiviral vector expressing EGFP fluorescence and puromycin resistance whereas B35 cells were transduced with a lentivirus expressing DsRed-Express2 fluorescence and hygromycin resistance, followed by isolation of clonal lines for use in fusion. Cells were fused in suspension at 1:1 ratio between the two cell types for 3 minutes at 37°C by adding a solution of 50% (w/v) PEG 1500 in serum-free DMEM pre-warmed to 42°C to pelleted cells. After washing, cells were plated in ESC media for ESC-somatic fusion or EpiSC media for EpisSC-somatic fusion. Puromycin and hygromycin were added the next day to select for fused cells. Successful fusion was confirmed by their double fluorescence. For hybrid clones, individual double fluorescent and double antibiotic resistant colonies were picked manually 10-15 days post-fusion and expanded. Three hybrid clones from the E14-B35 fusion, E14-B35(clone1), E14-B35(clone2) and E14-B35(clone3), and two hybrid clones for the G14-B35 fusion, G14-B35(clone9) (aka Fc9) and G14-B35(clone11) (aka Fc11), were used in the study. For ESC(EsrrbKO)-B35 and ESC(EsrrbKO)-G14 fusions, G14 cells were the same as that used in G14-B35 fusion; ESC(EsrrbKO) cells were transduced with a lentivirus expressing mCherry fluorescence and hygromycin resistance whereas B35 cells were transduced with a lentivirus expressing EGFP fluorescence and puromycin resistance, followed by isolation of clonal lines for use in fusion.

Cells were plated on a 10 cm dish at a 1:1 ratio between the two cell types, and fused 3 hours later on the plate for 1 minute by adding to the aspirated plate a solution of 45.5% (w/v) PEG 1000 in serum-free DMEM pre-warmed to 42°C. After washing, cells were plated in ESC media overnight, and puromycin and hygromycin were added the next day to select for fused cells that were further confirmed by their double fluorescence.

### RNA-seq

RNA samples were isolated using the MagNA Pure RNA isolation procedure (Roche) or with the RNAeasy Micro Kit (Qiagen) following manufacturer’s recommendations. RNA-seq library preparation was performed according to Illumina TruSeq recommendations. Samples were run on the Illumina HiSeq 2500 sequencer following vendor’s instructions. RNA-seq data were aligned to an orthologous mouse-rat ORF database we had constructed and the expression levels of individual genes were quantified as transcripts per genome (TPG) following previously described method with the modification that the average transcript length was set at 1600 nt (Looney et al. 2014). In G14-B35 fusions, the expression threshold (in TPG) for calling occluded genes in B35 is ≤5 in unfused B35, ≥10 in unfused G14, and ≥10 fold higher expression from mouse than rat genome in unfused G14 and unfused B35 and also in G14-B35 hybrids. Only genes with ≥90% ORF sequence identity between mouse and rat orthologs were included in the analysis to minimize false orthology.

### DNase-seq

DNase-seq was adapted from published protocol (Song and Crawford 2010). Cell lines were trypsinized and washed, followed by lysis with 0.025%-0.05% NP-40 based on cell type optimization. DNA was digested with DNase I at concentrations ranging from 0.4-2 U/ul. Digested DNA was visualized on a CHEF gel, and optimally digested samples were used for library preparation. Library preparation was performed according to Illumina recommendations. For data analysis, reads were mapped to a fusion-specific reference genome using Bowtie2 with default parameters. The fusion-specific reference genome was derived from MM10 mouse genome data and RN6 rat genome data downloaded from UCSC Genome Browser. Post-alignment SAM files were separated into species-specific SAM files and converted to BAM files. Duplicated reads were removed using Picard tools. MACS2 was used for Peak calling, with parameters --nomodel --shift --75 --extsize 150, and bedgraphs were normalized based on average peak heights between 10 and 60 percentiles, and then normalized with average peak heights of housekeeping genes. Normalized bedgraph files were converted to bigwig format and plotted on IGV. Average plots near the TSS of occluded genes were generated using normalized bedgraph files.

### Lentivirus transduction

Lentiviral vectors were constructed and packaged into virus by VectorBuilder. Vendor’s vector IDs for these vectors are listed below and can be used to retrieve detailed vector information from the vendor’s website. EGFP and puromycin resistance vector: VB150915-10026; mCherry and hygromycin resistance vector: VB150925-10020; Esrrb vector: VB180510-1202zrv; Klf4 vector: VB181219-1169xkg; mCherry control vector: VB160109-10005. Viral transduction of cells was performed according to vendor’s instructions.

### BAC modification and introduction into cells by piggyBac transposition

A BAC containing the entire rat Esrrb gene was purchased from the Children’s Hospital Oakland Research Institute BACPAC Resources Center (cat# CH230-121K10). It was modified by VectorBuilder to introduce into the vector backbone two piggyBAC inverted terminal repeats (ITRs) flanking the insert as well the puromycin resistance gene. The modified BAC along with pCMV-hyBase, a plasmid expressing the piggyBac transposase as published previously (Yusa et al. 2011), was electroporated into G14 and E14 cells, or chemically transfected into SiA cells. Puromycin was added 2 days later to select for cells that have stably incorporated the BAC via piggyBac transposition. For G14, total RNA and genomic DNA were harvested 10 days after transfection for RT-PCR and genomic PCR respectively. Primers for both RT-PCR and genomic PCR amplification of Esrrb are as follows: ATTCGGAGAACAGCCCCTAC, ACACAAGCTCCCGATCTGC. These primers are designed to amplify Esrrb from both mouse and rat copies. For RT-PCR, the product size is 189 bp for both mouse and rat Esrrb. For genomic PCR, the product size is 620 and 574 bp for mouse and rat Esrrb, respectively. RT-PCR product is sequenced using the following primers: GAGAACAGCCCCTACCTG, CAAGCTCCCGATCTGCCA. For E14, after cells with stably integrated rat Esrrb BAC have been selected by puromycin, cells were differentiated into EpiSC-like cells by culturing them in EpiSC media for 16 days before. Total RNA and genomic DNA were then harvested for RT-PCR and genomic PCR respectively. Primers for genomic PCR amplification of Esrrb were the same as those used on G14 transfected with rat Esrrb BAC. RT-PCR primers for various genes are as follows: Mouse-specific Esrrb: AGGTAGAGAAGGAAGAGTTTATGATCCTCAAG, GTCCGTCCGTCCATGTGCT; rat-specific Esrrb: GGAGAAGGAAGAGTTTGTGATGCTCAAA, GTCCATCCGTCTGCATGCG; mouse Pou5f1: GTTCAGCCAGACCACCATCT, ACTCCACCTCACACGGTTCT; mouse Klf4: AAGAACAGCCACCCACACTT, GGTAAGGTTTCTCGCCTGTG; mouse Fgf5: GGGATTGTAGGAATACGAGGAGTT, TGGCACTTGCATGGAGTTT; mouse Actb: CGCAGCCACTGTCGAGTC, ACCCACATAGGAGTCCTTCTGAC.

### Bisulfite sequencing

Bisulfite sequencing was performed using EpiTech Bisulfite Kit (QIAGEN) following vendor’s instructions. PCR primers to amplify Pou5f1 distal enhancer region are as follow: Mouse-specifc primers: AGGTTAGGGTATATTTGTTTTAAGTTAGTTTTAAGAAG, CTCCCAATTTCTATACATTCATTATAAAACAATACCATAA; rat-specific primers: TAAGGTTATATAGGGAGTTTTTGAATGAAATATAAATAAATAAA, ACCCAATTCCCAAAACAAACACAAACTTCC. PCR products were cloned into plasmid using TOPO TA Cloning Kit (Invitrogen) and individual clones were picked for Sanger sequencing of the inserts.

### ChIP-seq data analysis

ChIP-seq data for Esrrb and Pou5f1 in mouse ESCs were retrieved from Gene Expression Omnibus (GEO). Esrrb ChIP-seq data were from the study by Chen et al. (Chen et al. 2008) in GEO series GSE11431, in which ES_Esrrb (GSM288355) data were used for the analysis with ES_GFP (GSM288358) data as input control for peak calling by MACS2. Pou5f1 ChIP-seq data were from the study by Marson et al. (Marson et al. 2008) in GEO series GSE11724. Oct4_mES_rep1 (GSM307137) data were used for data analysis with WCE_mES_rep1 (GSM307155) and WCE_mES_rep2 (GSM307154) combined data as input control for peak calling by MACS2. Data analysis was done by the same pipeline as DNase-seq data analysis with --nomodel --extsize 300 as parameters for MACS2 peak calling.

## ACKNOWLEDGMENTS

This work was funded in part by the Chicago Biomedical Consortium with support from the Searle Funds at The Chicago Community Trust and by the Frontier Explorer Foundation. We thank Drs. Austin Smith and Graziano Martello for generously providing the Esrrb knockout ES cells.

## AUTHOR CONTRIBUTIONS

Conceptualization and methodology: K.M.F. and B.T.L.; Investigation, formal analysis and visualization: K.M.F., J.H.L., L.Z., C.J.F., B.W. and J.G.; Software: J.H.L., S.W.B. and T.J.L.; Resources: A.P.X. and G.F.; Writing: K.M.F. and B.T.L.; Supervision and funding acquisition: B.T.L.

