## Supplemental Figure S1 for "Capacity to erase gene occlusion is a defining feature distinguishing naive from primed pluripotency"

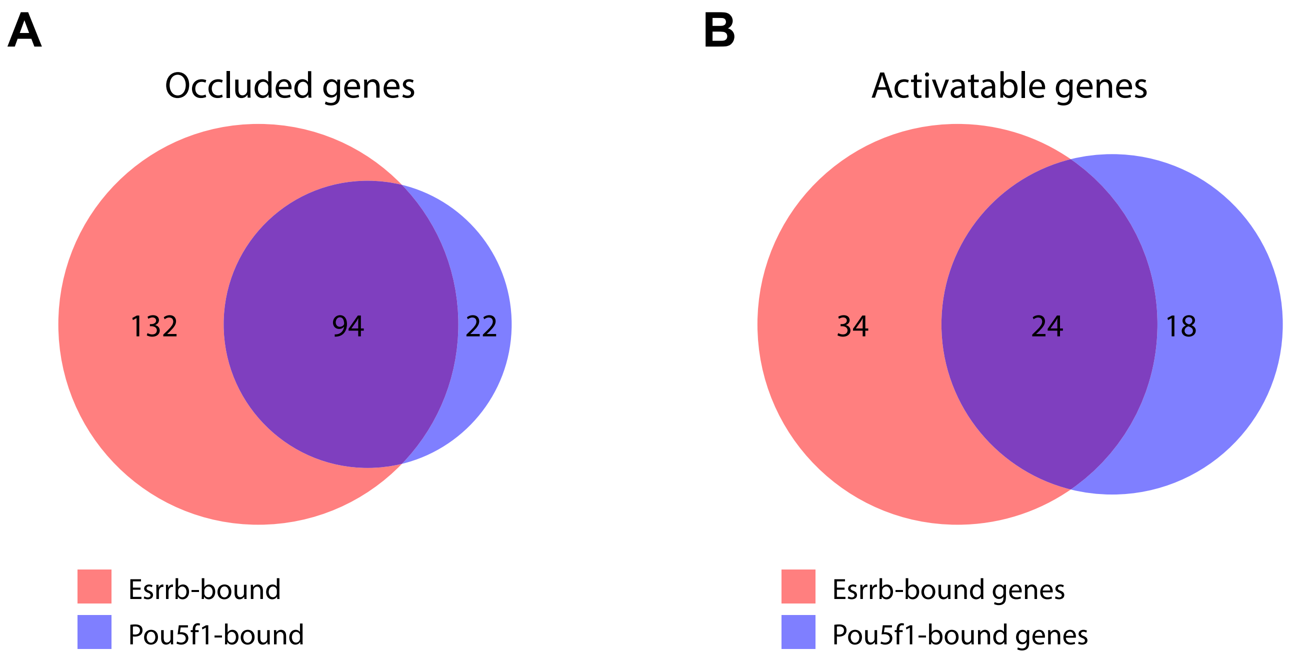


**Figure S1.** Venn diagram showing binding of Esrrb and Pou5f1 in occluded (A) or activatable (B) genes.
